# Safety Signals Enable Single-Episode Active Avoidance paradigm and Expose Threat Generalization in Tuberous Sclerosis Complex

**DOI:** 10.64898/2026.02.28.708704

**Authors:** Andrew Vincent Gallagher, Amber Victoria Wilson, Saheed Lawal, Harshil Sanghvi, Javed Iqbal, Matthew Dickinson, Brandon Li, Daniel Adolfo Llano, Prerana Shrestha

**Affiliations:** Graduate Program in Neuroscience, Stony Brook University, Stony Brook, NY 11794; Department of Neurobiology & Behavior, Stony Brook University, Stony Brook, NY 11794; Simons STEM Scholars Program, Stony Brook University, Stony Brook, NY 11794; Graduate Program in Computer Science, Stony Brook University, Stony Brook, NY 11794; Department of Molecular and Integrative Physiology, University of Illinois at Urbana-Champaign, IL 61801; Beckman Institute for Advanced Sciences and Technology, University of Illinois at Urbana-Champaign, IL 61801; Renaissance School of Medicine, Stony Brook University, NY 11794

## Abstract

Animals must flexibly discriminate between threat and non-threat to deploy adaptive defensive strategies. We introduce a single-episode differential signaled active avoidance (DSAA) paradigm that temporally dissociates acquisition, consolidation, and retrieval of instrumental avoidance memory. Interleaving a behaviorally noncontingent neutral cue (CS-) with a threat-predictive, behaviorally contingent cue (CS+) enhanced long-term memory without altering acquisition, demonstrating that differential contingency structure selectively reinforces memory consolidation rather than influencing learning performance. Naïve animals inferred contingencies within a single episode, and discrimination achieved during training predicted retrieval precision. However, discrimination operated within defined boundary conditions: overtraining or elevated threat intensity destabilized cue specificity and promoted persistent avoidance generalization. Under high-threat conditions, freezing and shuttling co-emerged as complementary defensive responses, indicating a shift from precise cue-based encoding to a generalized defensive state. Remote retrieval recruited oxytocin receptor-expressing cells in the medial prefrontal cortex and activated mTORC1-dependent translational signaling, implicating protein synthesis in maintenance of discriminative avoidance memory. In a Tuberous Sclerosis Complex model with Tsc2 haploinsufficiency in oxytocin-responsive cells, males displayed intact acquisition but generalized avoidance at both recent and remote time points, a deficit not rescued by additional training. These findings identify oxytocin-modulated translational control as a molecular gate stabilizing threat-safety discrimination and show that disruption of this axis - by excessive threat or reduced Tsc2 gene dosage - biases memory toward pathological generalization, providing a mechanistic framework for safety-learning deficits in neurodevelopmental and anxiety-related disorders.

## Introduction

Animals navigating dynamic environments must continuously discriminate between threat and safety to deploy appropriate defensive strategies. Defensive behaviors span reactive responses such as freezing and passive avoidance, and proactive responses including active escape or avoidance^1,2^. Threat imminence theory posits that defensive reactions scale with proximity and controllability of danger, shifting from freezing to flight as threats become imminent^3^. Yet when threats are predictable and controllable, adaptive behavior requires more than reflexive responding: animals must flexibly suppress defensive reactions during non-threat cues while rapidly engaging goal-directed avoidance when danger-predictive cues arise^4,5^. Effective threat regulation therefore depends on precise cue discrimination - minimizing unnecessary defensive expenditure during safety to conserve metabolic resources, while enabling swift, strategic action when harm is signaled. Disruption of this discrimination process is a defining feature of anxiety disorders, post-traumatic stress disorder, and neurodevelopmental conditions marked by emotional dysregulation, where exaggerated threat generalization and impaired safety signaling drive persistent maladaptive behavior^6–10^.

Despite its ethological relevance, instrumental active avoidance remains comparatively understudied at the mechanistic level. Most existing paradigms require multi-day training and performance criteria before memory assessment^11–13^, resulting in overlap between acquisition, consolidation, and retrieval phases. This temporal entanglement has limited circuit-level dissection of avoidance memory relative to single-episode Pavlovian threat conditioning, which has enabled precise interrogation of consolidation-dependent plasticity^14,15^. Yet in real-world settings, a single aversive encounter can produce enduring avoidance behaviors. To model this learning dynamic, we developed a single-episode differential signaled active avoidance (DSAA) paradigm in which a threat-predictive cue (CS+) requiring instrumental shuttling is interleaved with a neutral safety cue (CS-). This structure permits temporal separation of memory phases while introducing an explicit safety signal that constrains threat prediction. The DSAA paradigm builds upon the traditional signaled active avoidance framework, historically influenced by the two-factor “fear-avoidance” theory. This theory posits that reduction of fear elicited by the Pavlovian conditioned stimulus (CS) serves as the primary reinforcer driving the initial avoidance learning^5,16–18^.

Here we show that incorporation of a safety cue enables robust single-episode consolidation of active avoidance memory without altering acquisition. Discrimination achieved during training predicts long-term memory precision, and naïve animals rapidly infer contingencies without prior instruction. However, stimulus specificity operates within a defined dynamic range: excessive reinforcement or elevated threat intensity destabilizes discrimination and promotes persistent threat generalization. At the circuit level, remote retrieval recruits oxytocin receptor–expressing cells in the medial prefrontal cortex and engages mTORC1-dependent translational signaling, implicating protein synthesis in discriminative avoidance memory. Using a Tuberous Sclerosis Complex (TSC) mouse model with Tsc2 haploinsufficiency in oxytocin-responsive cells^19^, we identify a male-specific impairment in safety cue-dependent discrimination characterized by generalized avoidance despite intact acquisition. Together, these findings establish DSAA as a sensitive single-episode assay for dissecting the neural mechanisms of active avoidance and reveal oxytocin-modulated translational control as a molecular gate that stabilizes threat–safety discrimination and constrains pathological generalization.

## Results

### Adding a Safety Cue Improves Consolidation and Expression of Signaled Active Avoidance

Consistent with our previously established threat discrimination protocol^20^, we selected 7.5 kHz and 3 kHz tone frequencies to serve as the CS+ and CS-, respectively, in the DSAA paradigm. To exclude the possibility that behavioral differences were driven by unequal tone audibility, we assessed auditory brainstem responses (ABRs) in male and female C57BL/6 mice. Under anesthesia, hearing sensitivity was quantified by recording brainstem-evoked potentials in response to tone bursts presented at progressively increasing sound intensities, allowing determination of auditory thresholds for each frequency. Both sexes exhibited comparable hearing ranges, with higher thresholds for the 3 kHz CS- tone (∼70 dB) than for the 7.5 kHz CS+ tone (∼45 dB) (**Fig. 1A**). Accordingly, we adjusted the stimulus amplitudes such that CS+ and CS- were presented at 60 dB and 85 dB, respectively, ensuring that both cues were well above threshold and perceived at comparable salience.

**Figure 1.**
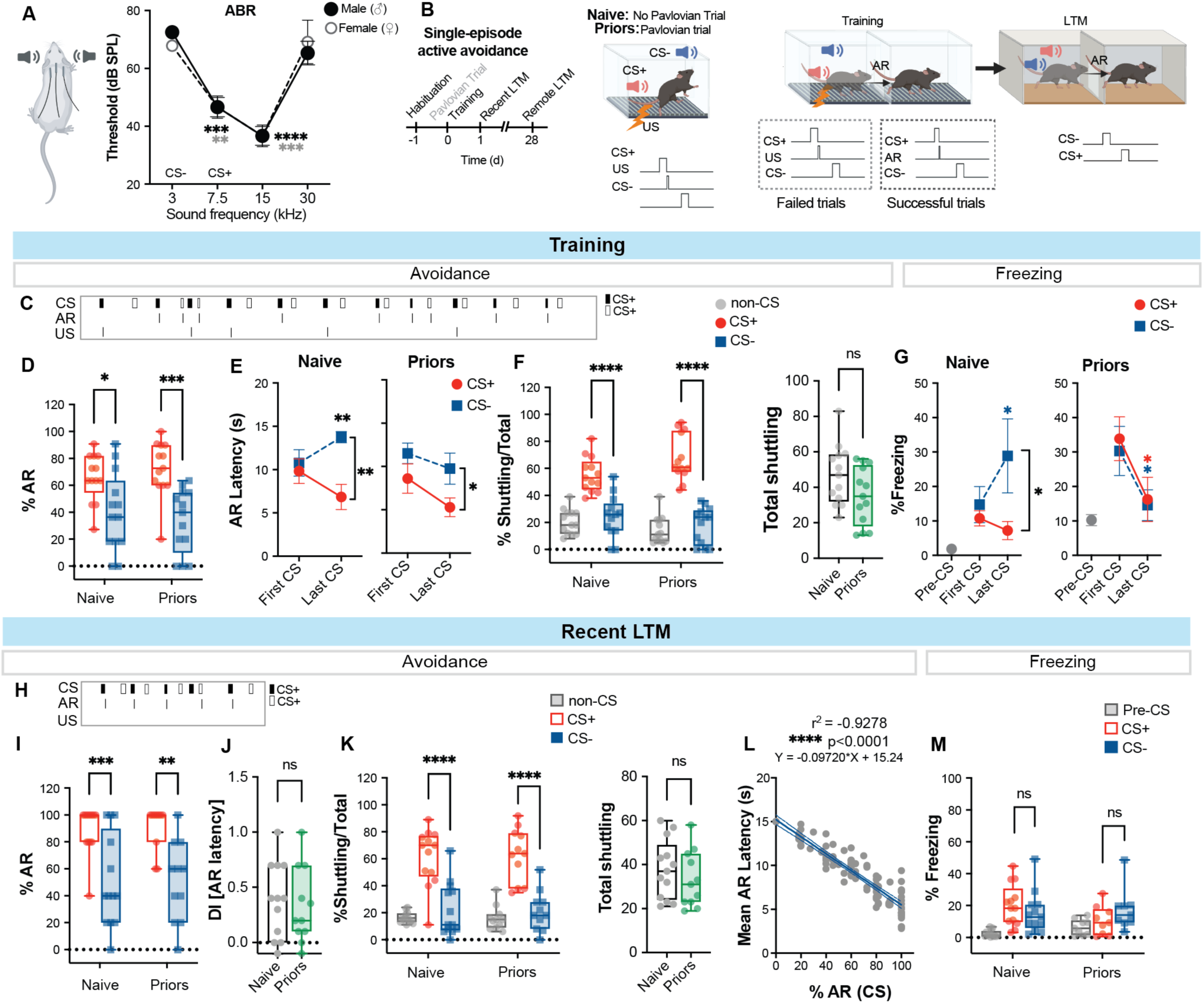
A) Auditory brainstem responses (ABRs) recorded from male and female C57BL/6J mice across a range of sound frequencies (3, 7.5, 15, and 30 kHz). **B)** Schematic of the single-episode differential signaled active avoidance (DSAA) paradigm. Mice undergo a single training session, preceded by a Pavlovian conditioning trial in the priors group, followed by longitudinal assessment of recent and remote long-term memory (LTM) at 24 hours and 28 days post-training. Failed trials are defined as trials in which mice do not shuttle during the CS+ and consequently receive the footshock unconditioned stimulus (US). Successful trials are those in which mice execute an avoidance response (AR) during the CS+, thereby preventing shock delivery. The neutral safety cue (CS-) carries no behavioral contingency. **C)** Representative event log illustrating the temporal occurrence of conditioned stimuli (CS+ and CS-), avoidance responses (AR), and unconditioned stimuli (US) for a single mouse during DSAA training. **D)** Box-and-whisker plots showing percent avoidance responses to CS+ and CS− during DSAA training in naïve and priors groups. **E)** Learning curves depicting changes in avoidance latency from the first to the last CS presentation for each cue (CS+ and CS−) in naïve (left) and priors (right) groups. **F)** Left: Distribution of shuttling behavior (expressed as percent of total shuttling) across CS+, CS−, and non-CS epochs. Right: Total shuttling during training in naïve and priors groups. **G)** Percent freezing during DSAA training measured during Pre-CS, first CS, and last CS epochs in naïve and priors groups. **H)** Representative event log illustrating the temporal occurrence of conditioned stimuli (CS+ and CS-), avoidance responses (AR), and unconditioned stimuli (US) for a single mouse during DSAA recent LTM test. **I)** Left: Cue-evoked avoidance responses during the recent LTM test. Right: Discrimination index (DI) based on avoidance latency for naïve and priors groups. **J)** Left: Shuttling distribution across CS+, CS−, and non-CS epochs during recent LTM testing. Right: Total shuttling during recent LTM in naïve and priors groups. **K)** Pearson correlation analysis demonstrating a strong positive correlation between mean avoidance latency and percent avoidance responses (training and LTM data pooled across CS+ and CS−; r² = 0.9278, ****p < 0.0001). **L)** Percent freezing during the recent LTM test in naïve and priors groups. **Statistical analyses:** A, D, F) RM Two-way ANOVA with Bonferroni post hoc test; C, E (left), G, I (left), K) Two-way ANOVA with Bonferroni post hoc test; E (right), H, I (right) Unpaired t-test. **Sample size**: n = 6–13 mice per group. *p < 0.05, **p < 0.01, ***p < 0.001, ****p < 0.0001; ns, not significant.

Conventional single-tone signaled active avoidance (SAA) is characterized by relatively slow acquisition and typically requires multiple days of training. For this paradigm, we presented mice with 11 presentations of CS+ (7.5 kHz pulse for 15s) that predicts an impending footshock (0.2 mA) and must shuttle during the tone to avoid shock delivery. No additional auditory cues are provided. To test whether incorporating a neutral auditory tone enhances learning or memory retention, we implemented a DSAA protocol in which the CS+ presentations were interleaved with CS- trials (3 kHz pure tone for 15s) (**Supplementary Fig. 1A**). During training, SAA and DSAA mice performed comparably, with no significant differences in percent avoidance or avoidance latency (**Supplementary Fig. 1B, C**). Total shuttling and percent freezing were also similar between groups (**Supplementary Fig. 1D, E**), indicating equivalent baseline locomotor activity and defensive responses. Despite similar acquisition rates, recent long-term memory (LTM) performance diverged. DSAA-trained mice exhibited superior memory retrieval, marked by higher avoidance rates, shorter avoidance latencies, increased shuttling, and reduced freezing relative to SAA-trained mice, as well as performance during the first day (**Supplementary Fig. 1F–I**). These findings suggest that inclusion of a safety cue enhances consolidation of CS+-evoked active avoidance.

### Contingency Learning in Differential SAA Is Robust in the Absence of Prior Information

To test whether priors influence the DSAA performance, we implemented a strategy in which mice either received explicit tone-shock contingency information (“priors”) or remained naïve. The priors group received a Pavlovian conditioning trial before the DSAA training. The Pavlovian trial consisted of a single presentation of the CS+ (7.5 kHz pulsed tone, 15 s) immediately followed by an unavoidable footshock (US; 0.2 mA, 0.5 s). Within the same trial, the CS- (3 kHz pure tone, 15 s) was presented after a 90s interval. Naïve mice did not receive this Pavlovian pre-exposure. (**Fig. 1B**). Long-term memory was assessed at two time points: 24 h after training and again 28 days later in a novel context, to test whether the auditory CS could elicit an instrumental avoidance response in the absence of reinforcement.

On the training day, both naïve and priors groups exhibited a significantly higher proportion of avoidance responses during CS+ presentations compared to CS- (**Fig. 1C, D**). Analysis of avoidance latencies across training revealed similar learning dynamics between groups. In the naïve group, avoidance latency decreased across CS+ trials but increased across CS- trials, resulting in a marked enhancement of cue discrimination. The priors group showed modest but significant changes in avoidance latency between early and late trials (**Supplementary Fig. 2A, Fig. 1E)**. Total shuttling activity and the distribution of shuttling across non-CS, CS+, and CS- epochs did not differ between groups. In both groups, shuttling was highest during CS+ presentations, whereas shuttling during CS- and non-CS epochs remained comparably low (**Fig. 1F**), indicating that group differences in discrimination were not driven by differences in overall locomotor activity. Because passive and active defensive responses are often inversely related, we next examined freezing behavior across training. In the naïve group, freezing increased from first to last CS- trials but remained stable across CS+ trials. In contrast, the priors group exhibited a steady decline in freezing across training for both CS+ and CS- (**Fig. 1G**).

During the recent long-term memory (LTM) test conducted in a novel context - distinguished by olfactory, visual, and somatosensory cues - both naïve and priors groups exhibited a significantly higher proportion of avoidance responses during CS+ relative to CS-, despite the absence of reinforcement (**Fig. 1H, I**). This produced a robust discrimination index based on avoidance latency between CS+ and CS−, alongside a comparable distribution of shuttling across CS and non-CS epochs in both groups (**Fig. 1J**). Moreover, the percentage of avoidance responses was strongly correlated with mean avoidance latency for both CS types (**Fig. 1K**), indicating that across training and memory retention testing, animals executed the instrumental avoidance response with progressively shorter latencies as cue-outcome associations were consolidated. In addition, discrimination indices measured during training were strongly positively correlated with those observed during the recent long-term memory test (**Supplementary Fig. 2B**), indicating that successful avoidance performance during memory retrieval directly reflected learning achieved during the training episode. Freezing responses were minimal in both naïve and priors groups, and no clear discrimination was observed between CS+ and CS− based on freezing behavior (**Fig. 1L**).

### Overtraining Impairs Stimulus Discrimination During Remote Retrieval of Cued DSAA Memory

We next asked whether additional training modifies the strength or specificity of DSAA memory at a remote time point. To address this, mice received two additional days of reinforced DSAA training in the same context (Context A) and were designated as “Overtrained.” Across these sessions, mice displayed increased avoidance responses and reduced response latencies to both the CS+ and CS- (**Supplementary Fig. 3A, B**), while maintaining comparable stimulus discrimination (**Supplementary Fig. 3C**). Although the proportional distribution of shuttling across CS+, CS-, and non-CS epochs remained stable, total shuttling increased on the second day, consistent with heightened defensive engagement and enhanced contextual responding (**Supplementary Fig. 3D, E**). Notably, mice exhibited greater freezing to the neutral cue (CS-) on the first training day; however, freezing responses were minimal on subsequent days, resulting in a reduced discrimination index (**Supplementary Fig. 3F, G**).

When tested for remote long-term memory on day 28 in a distinct context (context B), both naïve and Overtrained mice exhibited robust avoidance responses to the CS+ and retained discrimination between the two conditioned stimuli (**Fig. 2A**). However, Overtrained mice showed elevated avoidance responses and reduced latency to the CS-, resulting in a significant reduction in the discrimination index (**Fig. 2B, Supplementary Fig. 3A**). Overtraining also altered the temporal distribution of shuttling, yielding more similar levels of responding across CS+, CS-, and non-CS epochs, whereas this distribution remained preserved in naïve mice (**Fig. 2C**). Importantly, overtraining did not induce generalized hyperactivity, as total shuttling remained unchanged (**Fig. 2D**). Freezing responses remained minimal in both groups, and no CS-specific freezing discrimination was observed (**Fig. 2E, F**).

**Figure 2.**
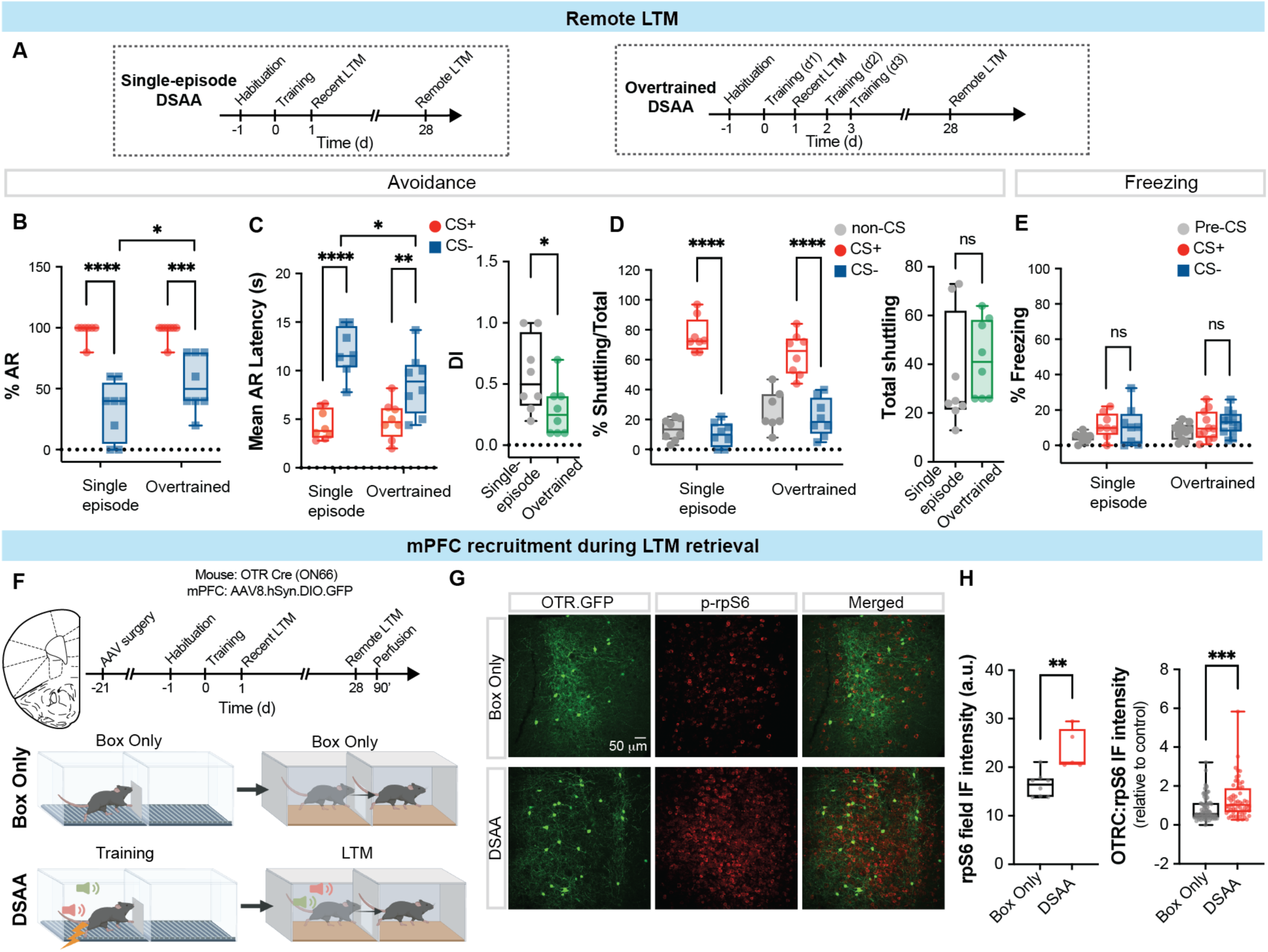
A) Schematic comparing the single-episode DSAA protocol with the overtrained DSAA protocol, which included two additional days of reinforced training. **B)** Box-and-whisker plots depicting percent avoidance responses to CS+ and CS− in single-episode versus overtrained groups. **C)** Left: Mean avoidance latencies to CS+ and CS− during remote LTM testing for single-episode and overtrained groups. Right: Discrimination index, showing a significant reduction in overtrained mice compared to single-episode controls. **D)** Left: Distribution of shuttling behavior (expressed as percentage of total shuttling) across CS+, CS−, and non-CS epochs for single-episode and overtrained groups. Right: Total shuttling during the remote LTM test. **E)** Percent freezing during pre-CS, CS+, and CS− epochs for single-episode and overtrained groups, demonstrating minimal freezing across conditions. **F)** Schematic illustrating labeling of oxytocin receptor–expressing cells (OTRCs) in the medial prefrontal cortex (mPFC), DSAA training, and longitudinal LTM testing. Tissue was collected 30 minutes after memory retrieval on day 28. Box-only controls were exposed to the training and testing contexts without cue or shock presentation. **G)** Representative images of phosphorylated ribosomal protein S6 (p-rpS6) expression in OTRCs from Box-only and DSAA-trained mice. Scale bar, 50 µm. **H)** Left: Quantification of p-rpS6 immunofluorescence (IF) field intensity. Right: Quantification of p-rpS6 IF within OTRCs, demonstrating increased activation in DSAA mice relative to Box-only controls. **Statistical analyses:** Panels B–E, two-way ANOVA with Bonferroni post hoc test; Panels C (right), D (right), and H, unpaired t-test. **Sample size:** Panels B-E, n = 6–8 mice per group; Panel H, n = 59–87 neurons from 3 mice per group. p < 0.05, p < 0.01, p < 0.001, p < 0.0001; ns, not significant.

The medial prefrontal cortex has been implicated in the expression of both passive and active avoidance behaviors^21–25^. Within this region, oxytocin receptor-expressing cells (OTRCs) are proposed to disinhibit layer II/III pyramidal neurons, thereby enhancing local network excitability and constraining anxiety- and stress-related responses^19,26,27^. Oxytocinergic neuromodulation of limbic–prefrontal circuits is thought to promote discrimination between stimuli that predict threat versus safety^28–30^. Consistent with this view, oxytocin signaling in the central amygdala has been shown to facilitate a transition from passive freezing to active escape in response to imminent but escapable threats, while attenuating reactivity to more distal threats^31^. However, whether oxytocin modulates prefrontal network activity during signaled active avoidance remains unknown.

To determine whether DSAA memory retrieval engages prefrontal OTRCs, we used OTR:Cre transgenic mice (ON66)^32^ and injected a cre-dependent viral vector expressing GFP into the prelimbic cortex (**Fig. 2G, H**). Brain tissue was collected 30 min after remote DSAA memory retrieval. Control mice expressing GFP in prelimbic OTRCs were exposed to Contexts A and B during sham training and retrieval in the absence of conditioned or unconditioned stimuli, and tissue was harvested 30 min after Context B exposure on day 28. We observed robust activation of mammalian target of rapamycin complex 1 (mTORC1) signaling in the prelimbic cortex during retrieval, as indicated by increased phosphorylation of ribosomal protein S6 (p-rpS6)^15,33^ (**Fig. 2I**). Notably, p-rpS6 immunofluorescence was significantly elevated in prelimbic OTRCs relative to controls, indicating the recruitment of oxytocin-responsive cells during remote DSAA memory retrieval.

### Co-emergence of avoidance and freezing as complementary defensive responses during high threat

To define the dynamic range of DSAA memory, we systematically varied the intensity of the footshock unconditioned stimulus (US), delivering 0.1 mA (weak), 0.2 mA (medium; standard DSAA), or 0.4 mA (strong) shocks to separate cohorts of mice while holding all other parameters constant (**Fig. 3A**). We used the naïve DSAA protocol, in which animals infer tone-shock and tone-behavior contingencies through Bayesian updating under initial uncertainty. During training, all groups exhibited robust avoidance responses to the threat-predictive cue (CS+). However, mice in the strong US group generalized avoidance to the neutral cue (CS-) (**Fig. 3B**). Avoidance latencies diverged appropriately from early to late trials across all three groups (**Fig. 3C**). The distribution of shuttling across CS+, CS-, and non-CS epochs remained stable, with a greater proportion of shuttling occurring during CS+ presentations (**Fig. 3D**), and total shuttling did not differ between groups (**Fig. 3E**). In contrast, freezing responses scaled with US intensity (**Fig. 3F**). The weak US group displayed minimal freezing to either cue. The medium US group showed a selective increase in freezing to the CS- during training, paralleling changes in avoidance latency. Strikingly, mice exposed to the strong US exhibited a pronounced escalation of freezing to both CS+ and CS-. Importantly, this loss of freezing discrimination occurred despite preserved discrimination in avoidance latency during training in the high-threat group.

**Figure 3.**
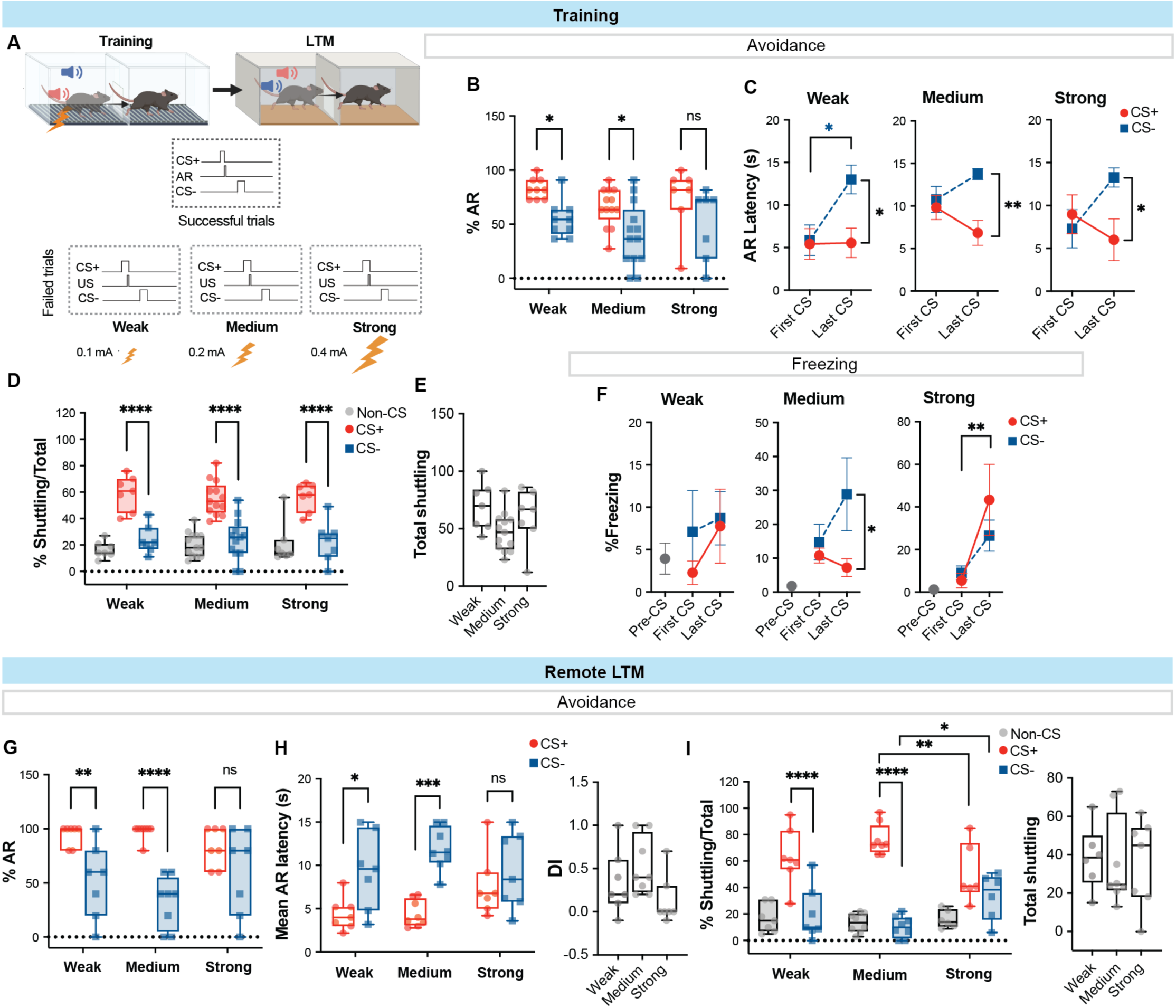
A) Schematic illustrating modulation of US intensity to define the dynamic range of DSAA learning and memory. Successful trials were defined identically across groups: shuttling during the CS+ prevented delivery of the footshock US. Failed trials (no shuttling during CS+) resulted in delivery of a weak (0.1 mA), medium (0.2 mA), or strong (0.4 mA) US, corresponding to group assignment. **B)** During DSAA training, differential avoidance responses were evident in the weak and medium US groups but were diminished in the strong US group. **C)** All three groups (weak, medium, and strong US) exhibited differential avoidance latency learning from the first to the last CS presentation. **D)** Distribution of shuttling behavior (expressed as percentage of total shuttling) across CS+, CS−, and non-CS epochs for weak, medium, and strong US groups during training. **E)** Total shuttling during DSAA training. **F)** Freezing responses during Pre-CS, first CS, and last CS epochs. Freezing to CS− escalated in both the medium and strong US groups, while freezing to CS+ also increased in the strong US group. **G)** During remote LTM testing, differential avoidance responses were maintained in the weak and medium US groups, whereas the strong US group exhibited memory generalization. **H)** Avoidance memory generalization was evident in the strong US group, while robust discrimination was preserved in the weak and medium US groups. **I)** Left: Distribution of shuttling behavior (percentage of total shuttling) across CS+, CS−, and non-CS epochs for weak, medium, and strong US groups during remote LTM. Right: Total shuttling across groups. **Statistical analyses**: Panels B, D, G, H, I: two-way ANOVA with Bonferroni post hoc test; Panels C, F: repeated-measures two-way ANOVA with Bonferroni post hoc test; Panels E, H (right), I (right): one-way ANOVA. **Sample siz**e: n = 6-13 mice per group. p < 0.05, p < 0.01, p < 0.001, p < 0.0001; ns, not significant.

We next evaluated recent and remote long-term memory (LTM) at 24 hours and 28 days after training, respectively. At the recent LTM time point, both the weak and strong US groups exhibited generalized avoidance responses and latencies to the neutral cue (CS−), whereas the medium US group maintained robust discrimination between CS+ and CS− (**Supplementary Fig. 4A, B**). The distribution of shuttling across CS+, CS−, and non-CS epochs diverged appropriately in the medium and strong US groups but converged in the weak US group, suggesting reduced memory retention in the weak group despite comparable total shuttling across all groups (**Supplementary Fig. 4C**). Cue-evoked freezing remained low in the weak and medium US groups but was elevated in the strong US group, without discrimination between CS+ and CS− (**Supplementary Fig. 4D**). At the remote LTM time point, the strong US group continued to display generalized avoidance responses and latencies, whereas the weak and medium US groups exhibited differential responding to CS+ versus CS− (**Fig. 3G, H**). Consistently, shuttling distributions diverged across CS+, CS−, and non-CS epochs in the weak and medium groups but converged in the strong US (high-threat) group. As a proportion of total shuttling, CS+ directed shuttling was reduced and CS− directed shuttling was increased in the strong US group relative to the medium US group (**Fig. 3I**). Freezing responses remained elevated to both CS+ and CS− in the high-threat group compared to the other groups (**Supplementary Fig. 3E**). Together, these results indicate that high threat during training drives persistent threat generalization in avoidance behavior while simultaneously elevating freezing to both cues, suggesting that animals in the high-threat condition recruit complementary defensive responses in which cue-evoked freezing precedes instrumental shuttling.

### Sex -specific Generalization of Cued Avoidance Memory in a TSC mouse model

We recently developed a mouse model of emotional dysregulation in Tuberous Sclerosis Complex (TSC), a neurodevelopmental disorder associated with a broad spectrum of neuropsychiatric symptoms, including anxiety^19^. In this model, constitutive deletion of one Tsc2 allele in oxytocin receptor–expressing (OTR⁺) cells (**Fig. 4A**) produces sex-specific vulnerability to social isolation in adulthood and disrupts innate avoidance behaviors across assays such as the open field, social approach, and marble burying tasks. We therefore asked whether learned avoidance behaviors are similarly affected in this TSC model. To address this, we used OTR.Tsc2^f/+^ mice and wild-type littermate controls, analyzing males and females separately given the distinct domains of innate avoidance deficits previously observed - males exhibiting impairments toward asocial targets and females toward social targets. All animals were reared under group-housed conditions to determine whether learned avoidance behaviors might be unmasked even in the absence of a chronic stressor such as social isolation, and whether they are more sensitive to Tsc2 haploinsufficiency than innate avoidance responses.

**Figure 4.**
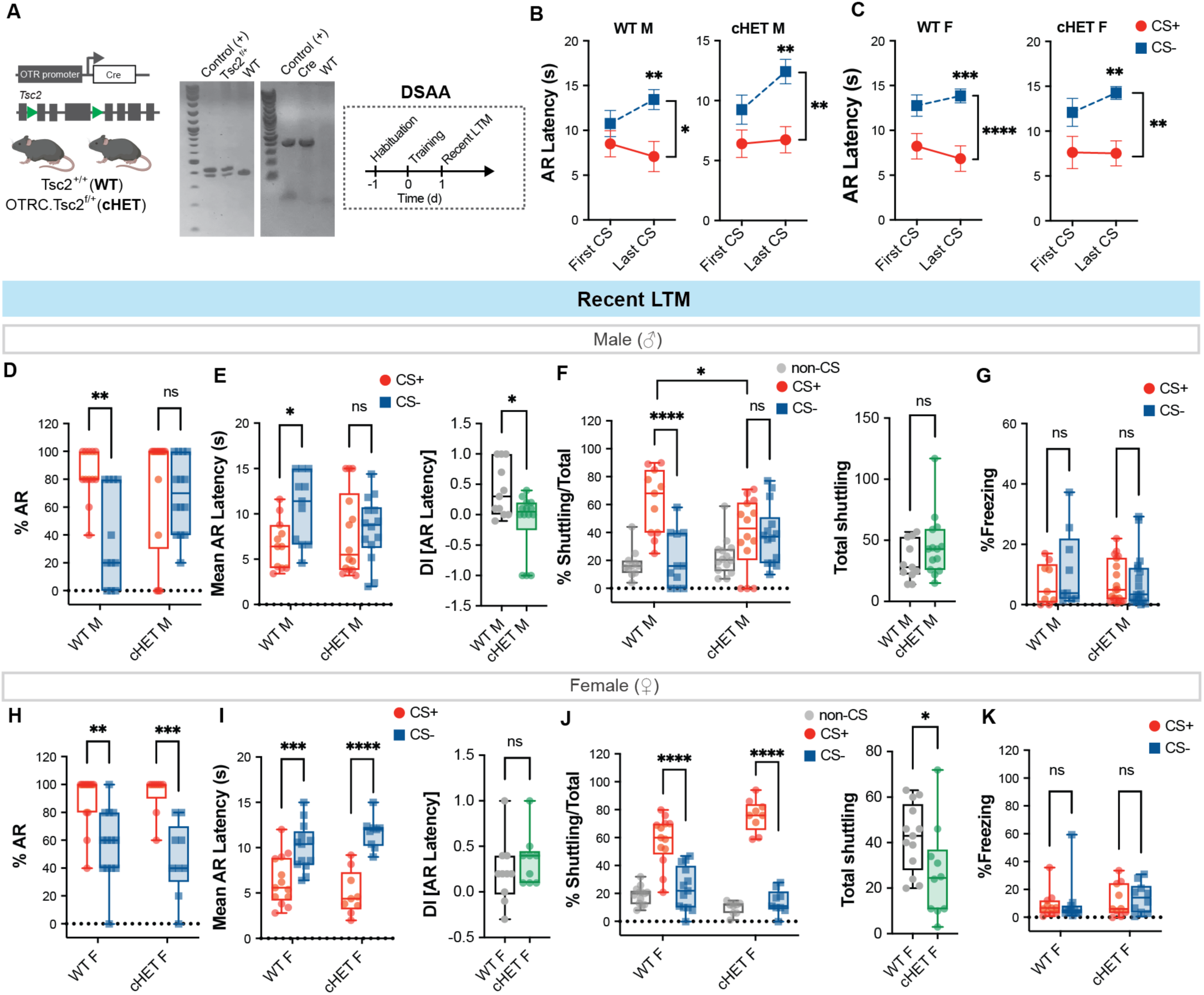
A) Breeding strategy used to generate OTR.Tsc2^f/+^ conditional heterozygous (cHET) mice and wildtype littermate controls. OTR-Cre transgenic mice were crossed with Tsc2^f/+^ mice. Genotyping with Cre-and Tsc2-specific primers identified offspring carrying both the Cre transgene and floxed Tsc2 allele. Both genotypes were subjected to single-episode DSAA training followed by long-term memory (LTM) testing. **B)** Learning curves depicting changes in avoidance latency from the first to the last CS presentation for each cue (CS+ and CS−) in males and females of both wildtype and cHET strains. **C)** During recent LTM testing, wildtype males exhibited robust differential avoidance to CS+ versus CS−, whereas cHET males failed to discriminate between cues. **D)** Left: Mean avoidance latencies during recent LTM in wildtype and cHET males. Right: Discrimination index (DI) for avoidance latency, demonstrating reduced discrimination in cHET males relative to wildtype controls. **E)** Left: Distribution of shuttling behavior (percentage of total shuttling) across CS+, CS−, and non-CS epochs during recent LTM in wildtype and cHET males. Right: Total shuttling during recent LTM. **F)** Cue-evoked freezing during recent LTM in male mice, which remained minimal across groups. **H)** Female wildtype and cHET mice both displayed robust differential avoidance responses during recent LTM testing. **H)** Left: Mean avoidance latencies for CS+ versus CS− in wildtype and cHET females. Right: Discrimination index (DI) for avoidance latency in female mice. **I)** Left: Distribution of shuttling behavior (percentage of total shuttling) across CS+, CS−, and non-CS epochs during recent LTM in wildtype and cHET females. Right: Total shuttling during recent LTM. **J)** Cue-evoked freezing during recent LTM in female mice, which remained minimal across genotypes. **Statistical analyses**: B) RM Two-way ANOVA with Bonferroni post-hoc test; C, D (left), E (left), F, G, H (left), I (left), J) Two-way ANOVA with Bonferroni post hoc test; D (right), E (right), H (right), I (right) Unpaired t-test. **Sample size:** n = 10–14 mice per group. *p < 0.05, **p < 0.01, ***p < 0.001, ****p < 0.0001; ns, not significant.

We subjected adult male and female mice to the naïve DSAA protocol using a medium-intensity US (0.2 mA), consisting of a single training episode followed by recent and remote LTM testing at 24 hours and 28 days post-training, respectively. During training, all groups exhibited comparable performance, characterized by robust acquisition of differential avoidance latencies, high levels of avoidance responding, and appropriate divergence in shuttling behavior during CS+ relative to CS− and non-CS epochs (**Fig. 4B-C, Supplementary Fig. 5A–D**). Notably, cue-evoked freezing strategies diverged by sex. Males progressively increased freezing to the neutral cue (CS−) across training, whereas females showed a reduction in freezing to the threat-predictive cue (CS+) over time (**Supplementary Fig. 5E, F**). These findings point to a sex-specific allocation of defensive strategies during reinforced learning: in males, the CS+ predominantly engages instrumental avoidance, whereas the neutral CS-, despite lacking behavioral contingency, appears to become incorporated into the threat context and elicits passive freezing, potentially through co-allocation within a shared temporal encoding window. Of note, we did not observe any genotype-effects during training.

We next assessed recent long-term memory (LTM) 24 hours after training. Compared to wildtype males, OTR.Tsc2^f/+^ cHET males exhibited generalized avoidance memory, reflected in both percent avoidance and avoidance latency (**Fig. 4D, E**). This generalization led to a significant reduction in the discrimination index in cHET males. The distribution of shuttling responses across CS+, CS−, and non-CS epochs also converged in cHET males, indicating impaired retention of cue-specific memory in the novel context, despite normal levels of total shuttling (**Fig. 4F**). Cue-evoked freezing remained minimal in both wildtype and cHET males (**Fig. 4G**). In contrast, female cHETs displayed robust differential avoidance memory, as evidenced by appropriate discrimination in both avoidance percentage and latency (**Fig. 4H, I**). The distribution of shuttling responses across CS+, CS-, and non-CS epochs diverged appropriately in both wildtype and cHET females; however, total shuttling was significantly reduced in cHET females (**Fig. 4J**). Freezing responses were minimal in both groups (**Fig. 4K**).

To determine whether the generalized avoidance observed in cHET males reflected inadequate discrimination learning after a single training episode, we extended training by two additional days and assessed remote LTM 28 days later. Additional training enhanced overall avoidance performance and reduced latencies across genotypes (**Supplementary Fig. 6A, B**). Wildtype males retained robust discrimination between CS+ and CS−, as evidenced by differential avoidance responses and latencies. In contrast, cHET males continued to generalize between the cues (**Supplementary Fig. 6C, D**). Consistent with this, the distribution of shuttling responses across CS+, CS−, and non-CS epochs remained convergent in cHET males but diverged appropriately in wildtype controls, despite comparable total shuttling between groups (**Supplementary Fig. 6E**). Freezing responses remained minimal in both groups (**Supplementary Fig. 6F**).

Together, these findings indicate that male OTR.Tsc2^f/+^ cHET mice exhibit a specific impairment in the consolidation and/or retrieval of associative cue-evoked avoidance memory, rather than a deficit in task acquisition or locomotor output. This behavioral phenotype is consistent with the reduction in nascent protein synthesis observed in medial prefrontal oxytocin receptor–expressing cells (OTRCs), a molecular process critical for long-term memory consolidation^19^. Moreover, the engagement of the mTORC1 pathway in prefrontal OTRCs during memory retrieval (**Fig. 2H**) suggests that sustained translational control within these cells supports persistent cellular plasticity required for accurate discrimination between threat-predictive and neutral cues.

## Discussion

Although conventional SAA and DSAA produced comparable acquisition during training, only DSAA strengthened recent retrieval, yielding higher avoidance rates, shorter response latencies, and reduced freezing. Discrimination during training strongly predicted discrimination at retrieval, and avoidance probability closely tracked latency reductions, indicating consolidation of a temporally precise cue–action association. Notably, this enhancement did not depend on prior Pavlovian conditioning. Naïve animals inferred contingencies within a single episode, suggesting that differential cue structure itself facilitates Bayesian updating under uncertainty. The neutral cue therefore appears to reduce ambiguity during encoding, sharpening consolidation of CS+ specific instrumental memory.

More broadly, incorporation of a neutral cue into the avoidance paradigm enhances consolidation and long-term expression of cue-specific memory without altering acquisition dynamics. Why the neutral safety cue facilitates consolidation remains an open question. One possibility is that it engages flexible internal models formed in the medial prefrontal cortex (mPFC), which link environmental stimuli to aversive outcomes - even when associations are indirect^25,34^. The inclusion of a neutral cue may enrich the representational structure of the learning environment, promoting more robust inferential encoding and stabilization of memory traces. Inferential memory representations are thought to emerge through multi-step encoding mechanisms involving recruitment and stabilization of mPFC neurons, particularly those projecting to the amygdala. This framework may explain why the CS- transiently evokes defensive freezing during training, which diminishes across sessions yet persists at remote time points in high-threat conditions. At the same time, differential avoidance memory remains highly sensitive to boundary conditions, including overtraining, heightened threat intensity, and genetic disruption of translational plasticity.

The single-episode nature of DSAA may also depend on species-specific defensive biases. Defensive strategy selection is influenced more strongly by body size than by sex or age: smaller animals preferentially adopt proactive avoidance, whereas larger animals tend toward reactive freezing^35^. Thus, the efficiency of single-episode DSAA may be particularly suited to mice, which are substantially smaller than Long-Evans rats^35^. In contrast, previous rat studies required prolonged cue durations (up to 240 s), extensive reinforcement (30 trials per session), and multi-day training to enhance avoidance. In our protocol, a modest 15 s CS+ presented only 11 times, interleaved with a temporally distinct CS-, was sufficient to generate robust long-term memory. Importantly, DSAA does not include an explicit feedback signal following the avoidance response (AR). Animals infer action value solely through omission of the footshock. Although the CS− appears after the AR, it is temporally separated by 20-90 s and therefore does not meet the criteria for feedback-based reinforcement. Under standard and overtrained conditions, neither CS+ nor CS− evokes substantial freezing during retrieval, indicating suppression of central amygdala–periaqueductal gray circuits in favor of proactive avoidance. Only high-threat conditions produce co-expression of freezing and avoidance, accompanied by loss of cue discrimination. Manipulating shock intensity further revealed that discrimination operates within an optimal threat range. Standard (0.2 mA) footshock produced the sharpest divergence in avoidance responses, whereas both weaker and stronger shocks reduced response precision. These findings are consistent with Mowrer’s two-factor theory^17,36^: avoidance and freezing are goal-directed behaviors serving to reduce fear, but their balance depends on arousal level.

At the circuit level, remote DSAA retrieval robustly engaged prefrontal oxytocin receptor-expressing cells (OTRCs) and activated mTORC1-dependent translational signaling, as indicated by increased phosphorylation of ribosomal protein S6^37^. These findings suggest that retrieval of discriminative avoidance memory recruits protein synthesis-dependent plasticity within oxytocin-sensitive prefrontal ensembles. Retrieval is therefore not a passive readout but requires sustained translational control to stabilize or reactivate cue-specific neural representations. This aligns with models in which oxytocin signaling constrains anxiety-related responding and promotes discrimination between threat and safety within limbic–prefrontal circuits^28,30^. Consistent with this framework, Tsc2 haploinsufficiency in OTR-expressing cells produced a male-specific impairment in associative avoidance memory. Male OTR.Tsc2^f/+^ cHET mice displayed intact acquisition but persistent generalization at both recent and remote time points, a deficit not rescued by additional training and not attributable to altered locomotion or freezing. Given that TSC2 regulates both mTORC1- and PERK-eIF2α dependent translational control^19,38–40^, these findings implicate disrupted protein synthesis in the failure to consolidate or retrieve cue-specific memory. Together, the behavioral and molecular data support a model in which oxytocin-modulated translational control in the emotional memory network gates the balance between discrimination and generalization. When this regulatory axis is compromised - by excessive threat or impaired TSC signaling - precise avoidance encoding collapses into generalized defensive responding, providing a mechanistic framework for safety-learning deficits in neurodevelopmental and stress-related disorders.

## Methods

### A. Mice

Mice were provided with food and water ad libitum and were maintained in a 12h/12h light/dark cycle at Stony Brook University or University of Illinois at Urbana-Champaign at stable temperature (78oF) and humidity (40 to 50%). All mice were backcrossed to C57Bl/6J strain for at least 5 generations. Both male and female mice, aged 3-6 months, were used in all experiments. Floxed Tsc2 mice were kindly provided by Dr. Michael Gambello (Emory University). OTR Cre BAC transgenic mice (Founder line #ON66) were generated by GENSAT and kindly provided by Dr. Nathaniel Heintz (The Rockefeller University). Wildtype C57Bl/6J mice (stock #000664) were purchased from Jackson labs. All procedures involving the use of animals were performed in accordance with the guidelines of the National Institutes of Health and were approved by the University Animal Welfare Committee of Stony Brook University and University of Illinois at Urbana-Champaign.

### A. Genotyping

OTR-Cre mice were genotyped using the forward primer (5′-CCG GTG AAC GTG CAA AAC AGG CTC TA-3′) and reverse primer (5′-CTT CCA GGG CGC GAG TTG ATA GC-3′). PCR amplification was performed under the following thermocycler conditions: 95°C for 5 min; 95°C for 30 s, 63°C for 45 s, and 72°C for 45 s (30 cycles); followed by 72°C for 10 min and hold at 4°C. A 408 bp band, present in the positive control and absent in negative controls, was considered indicative of the Cre genotype. Tsc2 floxed mice were genotyped using a forward primer (5′-GCA GCA GGT CTG CAG TGA AT-3′), reverse primer 1 (5′-CCT CCT GCA TGG AGT TGA GT-3′), and reverse primer 2 (5′-CAG GCA TGT CTG GAG TCT TG-3′). PCR conditions were as follows: 94°C for 2 min 50 s; 94°C for 30 s, 65°C for 1 min, and 68°C for 1 min (10 cycles); followed by 94°C for 15 s, 60°C for 1 min, and 72°C for 1 min (28 cycles); with a final extension at 72°C for 7 min and hold at 4°C. A 390 bp band corresponded to the floxed allele, whereas a 320 bp band indicated the wild-type allele.

### B. Stereotaxic surgeries

Mice were anesthetized with the mixture of Ketamine (100 mg/kg) and Xylazine (10 mg/kg) in sterile saline (i.p. injection). Adequate sedation was determined by a lack of gentle toe pinch withdrawal reflex. A lubricant eye ointment (Genteal Tears; Alcon) was applied on the eyes to prevent ocular dehydration. Visual monitoring of respiration occurred periodically throughout surgery to confirm survival. Stereotaxic surgeries were carried out inside a Class II A2 biosafety cabinet (LabRepCo, # NU-677-500) on the Kopf stereotaxic instrument (Model #942), which was equipped with a Nanojector UMP3TA (WdI). Viral vectors were injected intracranially using 2.0 μl Neuros syringe (Hamilton, #65459-02) at 1 nl/s. Before removing the needle, an interval of 10 min was allotted for viral diffusion. Upon completion of intracranial injections, the incision of the scalp was closed using a tissue adhesive (3M Vetbond). Normothermia during surgery was maintained at 36.6oC by resting the mouse on a covered heating pad (7 cm X 7 cm) connected to a rodent warmer console (Stoelting, # 53800M). To prevent somatic dehydration, each mouse was subcutaneously injected with 500 μl of sterile saline before surgery, and further provided with hydrogel (Clear H2O, # 70-01-5022) in the recovery cage. General monitoring and analgesia using 3 mg/kg subcutaneous Ketoprofen (Zoetis) were provided up to three days post-surgery. To label Oxytocin receptor-expressing neurons in the medial prefrontal cortex, 200 nl of AAV8.hSyn.DIO.GFP (1.00 X 10^13^ GC/ml) was injected into the stereotaxic coordinates ([-2.00 mm anterior posterior (AP), +/-0.25 mm mediolateral (ML) and -1.60 mm dorsoventral (DV)] of either OTR Cre+/- wildtype mice. AAV8-hSyn-DIO-EGFP was a gift from Bryan Roth (Addgene plasmid # 50457 ; http://n2t.net/addgene:50457 ; RRID:Addgene_50457).

### C. Auditory brainstem responses

Wildtype C57Bl/6J male and female mice were anesthetized with intraperitoneal ketamine (100 mg/kg), xylazine (6 mg/kg), and acepromazine (3 mg/kg). After achieving surgical anesthesia, subdermal electrodes (active, recording, and ground) were placed at the vertex, behind the left ear, and behind the right ear, respectively. Tone bursts (5 ms duration; 1 ms rise/fall, 3 ms plateau) were presented at 21 Hz. A 3 kHz stimulus was delivered free field via an AS03608AS-R speaker (PUI Audio) positioned 5 cm ipsilateral to the recording ear, while 7.5, 15, and 30 kHz stimuli were delivered via a TDT ES1 speaker positioned 5 cm from the interaural midline. Speakers were driven by TDT System 3 hardware with RPvdsEx software and calibrated using a Svantek 977 Sound and Vibration Level Meter and Larson Davis CAL200 calibrator. Auditory brainstem responses (ABRs) were acquired using TDT RA4PA Medusa preamps, bandpass filtered (500–3000 Hz), and averaged over 512 sweeps within a 30 ms window. Thresholds were determined using a bracketing approach from 90 to 10 dB SPL. Coarse 10 dB decrements identified a response bracket, followed by 2 dB steps to define threshold as the lowest level producing identifiable waves I–V (0–10 ms), ≥100 nV amplitude, consistent morphology at +2 dB, and absence at −2 dB.

### D. Behavior

All behavior sessions were conducted during the light cycle (7:00 AM – 7:00 PM EST). Both male and female mice were included in behavior experiments. Behavior experiment data were collected by experimenters that were blind to the experimental conditions and genotype. Following weaning, mice were group-housed with up to three sex-matched littermates in standard cages (8 in X 13 in X 5 in) fitted with corn cob bedding and Enviro-dri enrichment. Acclimation of the mice in the behavior room lasted at least 30 min prior to testing. A white noise machine (Yogasleep Dohm) was used to blunt any environmental noise that may induce confounding anxiety. All items in the behavior equipment were cleaned with 30% ethanol unless otherwise stated.

#### Data acquisition

Behavioral experiments were performed using the Habitest Linc System (H01-01 Power Base and H02-08 Linc Boxes; Harvard Apparatus) controlled by Graphic State 4 software (Coulbourn Instruments), which regulated delivery of footshock, light stimuli, and auditory tones. Graphic State 4 also recorded shuttling behavior throughout each session. Freezing during DSAA training and memory retrieval was quantified using FreezeFrame 4 software (Actimetrics), which captured video and automatically scored freezing responses. Two distinct environmental contexts (Contexts A and B) were used for the SAA and DSAA paradigms. Context A consisted of a chamber (29 cm W × 23 cm D × 20 cm H) equipped with a white house light and metal shock grid floor, cleaned with 30% ethanol, and lacking striped wall panels. Context B was a chamber of similar dimensions but contained infrared illumination, striped wall panels, a white plexiglass platform floor, vanilla-scented paper bedding, and was cleaned with 30% isopropanol to provide distinct visual, tactile, and olfactory cues. Chambers were housed within a sound-attenuating isolation cubicle (30W × 19H × 18D; H10-24 and H10-24A, Coulbourn Instruments). Footshocks were delivered via a Precision Animal Shocker (H13-15, Coulbourn Instruments), and auditory stimuli were presented programmatically using an H12-01M speaker module coupled to an H12-07 seven-tone audio cue.

#### Single-tone signaled active avoidance

Training: Mice were acclimated to Context A for 120 s before training commenced. Training consisted of 11 CS+ presentations (7.5 kHz pulsed tone; 0.2 s on, 0.2 s off) delivered with a variable inter-trial interval (ITI; 20–90 s, mean 45 s). Avoidance was defined as a shuttle response during the CS+, which immediately terminated the tone and prevented footshock delivery. Shuttling is the movement from one chamber to the adjacent chamber through an unobstructed doorway separating the two chambers. The response was not direction- or chamber-specific but rather constituted the instrumental action of crossing between compartments. Failure to shuttle within the 15 s CS+ resulted in delivery of a 0.2 mA footshock (US) for 0.5 s. Shuttling during the shock period was classified as an escape response.

Recent Long-term memory (LTM) test: Twenty-four hours after training, mice were acclimated to Context B for 120 s prior to testing. They then received 5 CS+ presentations with a variable ITI (20–90 s, mean 45 s). Context B was defined by a white Plexiglas floor, striped walls, infrared illumination, vanilla-scented bedding, and 30% isopropanol as the olfactory cue. Trials were non-reinforced. Avoidance was measured as shuttling during the CS+, which terminated the tone. No footshock was delivered during the LTM test, even if the animal failed to avoid.

#### Pavlovian trial

The Pavlovian conditioning trial consisted of an unavoidable footshock delivered immediately following the CS+ in Context A. The CS+ was a 7.5 kHz pulsed tone (0.2 s on, 0.2 s off) presented for 15 s at 40% amplitude (approximately 60 dB) through a speaker mounted on the far wall of each chamber. Immediately after CS+ termination, a 0.2 mA steady footshock was delivered for 0.5 s. The CS- was a 3 kHz continuous tone presented at 60% amplitude (approximately 80 dB) for 15 s. The CS- was delivered 2 minutes after the unconditioned stimulus (US).

#### Differential signaled active avoidance (DSAA)

<u>Training</u>: Mice were acclimated to Context A for 120 s before training began. Training consisted of 11 CS+ presentations (7.5 kHz pulsed tone; 0.2 s on, 0.2 s off; 60 dB; ≤15 s duration) delivered with a variable inter-trial interval (ITI; 20–90 s, mean 45 s). Avoidance was defined as a shuttle response during the CS+, which immediately terminated the tone and prevented footshock delivery. Shuttling was defined as movement from one chamber to the adjacent chamber through an unobstructed doorway separating the two compartments. This response was not direction- or chamber-specific, but rather represented the instrumental act of crossing between compartments. Failure to shuttle within the 15 s CS+ resulted in delivery of a 0.2 mA footshock (US) for 0.5 s. Shuttling during the shock period was classified as an escape response. In addition, 11 CS− presentations (3 kHz continuous tone; 80 dB; ≤15 s duration) were interleaved with CS+ trials. The CS− was not reinforced. Shuttling during the CS− was scored as a CS−-evoked avoidance response.

<u>Recent LTM test</u>: Twenty-four hours after training, mice were acclimated to Context B for 120 s prior to testing. They then received 5 CS+ presentations with a variable ITI (20–90 s, mean 45 s). All trials were non-reinforced. CS+-evoked avoidance was measured as shuttling during the CS+, which terminated the tone. No footshock was delivered during the LTM test, even if the animal failed to avoid. Five CS- presentations were similarly interleaved with CS+ trials. CS+ and CS- evoked avoidance responses were quantified as shuttling during their respective cue presentations, each of which terminated the tone.

<u>Remote LTM test</u>: Remote long-term memory (LTM) was assessed in Context B twenty-eight days after training. Testing parameters were identical to those used for the recent LTM assessment.

#### Unconditioned stimulus intensity adjustments during DSAA training

Mice were trained using the standard DSAA protocol, with the exception that the intensity of the unconditioned stimulus (US) was systematically varied. The standard condition employed a 0.2 mA continuous footshock (medium US). In separate groups, the US intensity was adjusted to 0.1 mA continuous footshock (weak US) or 0.4 mA continuous footshock (strong US).

#### Overtraining

Mice were trained using the standard DSAA protocol across three consecutive days. Each day, animals were acclimated to Context A for 120 s, followed by 11 interleaved presentations of CS+ and CS−. Only the CS+ maintained the tone–shock contingency, such that failure to shuttle during the CS+ resulted in footshock delivery. Remote long-term memory (LTM) was assessed 28 days after the final training session in Context B. The test consisted of five interleaved CS+ and CS− presentations delivered with a variable inter-trial interval (20–90 s; mean 45 s).

#### Data processing pipeline

To facilitate efficient and reproducible analysis of avoidance conditioning experiments, we developed a fully automated, Python-based computational pipeline for data extraction, behavioral metric computation, and visualization. The workflow integrates timestamped event logs from FreezeFrame (freezing during Pavlovian threat conditioning and differential signaled active avoidance) and GraphicState (threat-predictive conditioned stimulus [CS+], safety cue [CS-], and unconditioned stimulus [US] -evoked responses) to generate structured, analysis-ready datasets. Custom modules (freezeframe.py, graphicstate.py) implement preprocessing steps including column name standardization, epoch segmentation (pre-CS, CS+, CS- for freezing behaviors, and non-CS, CS+, CS- for shuttling responses), metadata integration, and handling of missing data. Task-specific extensions (dsaa_gs.py) were generated to classify avoidance trial outcomes (avoidance, escape, failure), compute spontaneous and CS+ and CS- evoked shuttling, and calculate latency to avoid using precisely parsed CS onset/offset timestamps. On the other hand, dsaa_ff.py was generated to quantify freezing percentages across defined epochs (pre-CS, CS+ and CS-). The pipeline uses an object-oriented modular design built around the FreezeFrame and GraphicState base classes, supporting reusability across experimental paradigms. Automated batch processing was implemented for high-throughput analysis of multiple datasets with minimal manual input. Comprehensive logging and error handling (via logging.info(), logging.warning(), try-except blocks) were also utilized to ensure data integrity and traceability. Dedicated plotting modules (fr_pav_heatmaps.py) were developed to generate standardized visualizations of freezing dynamics, allowing rapid and reproducible figure generation. Together, this custom pipeline provided a scalable and transparent framework for behavioral data analysis in differential signaled active avoidance paradigm.

### E. Histology

Mice were deeply anesthetized with isoflurane, and transcardially perfused with 0.1M PBS followed by 4% paraformaldehyde (EMS) in PBS. Brains were removed and postfixed in 4% PFA for 24h. The PFA is then replaced with 30% sucrose solution for another 24 h. 40 μm free-floating coronal brain sections containing PFC were collected using Leica vibratome (VT1000s) and stored in 1X PBS containing 0.05% sodium azide at 4oC. After blocking in 5% normal goat serum in 0.1 M PBS with 0.1% Triton X-100, brain sections were probed overnight with primary antibodies [chicken anti-EGFP (Abcam #ab13970) and rabbit anti-p-S6 (S235/6) (Cell Signaling #4898, Lot #21). After washing three times in 0.1M PBS, brain sections were incubated with Alexa Fluor conjugated secondary antibodies 1:200 (Thermo Fisher A32931 Lot # VK314825, A32740 Lot # WC316326) in blocking buffer for 1.5h at RT and mounted using Prolong Gold antifade mountant (Fisher # P36930).

### F. Imaging

Imaging data from immunohistochemistry experiments were acquired using a laser scanning microscopy (LSM) confocal microscope (Zeiss) with 20X objective lens (with 1X or 2X zoom) and z-stacks (approximately 6 optical sections with 0.563 μm step size) for three coronal sections per mouse from AP Bregma +1.98 mm to + 1.78 mm (n = 3 mice) were collected. Imaging data was analyzed with ImageJ using the Bio-Formats importer plugin. Maximum projection of the z-stacks was generated followed by manual outline of individual cells and mean fluorescence intensity measurements using the drawing and measure tools. Mean fluorescence intensity values for all cell measurements were normalized to the mean fluorescence intensity for controls. In the case of assessing mTORC1 pathway activation in mPFC, field fluorescence of the entire image area was measured, instead of outlining each cell. Each image was opened with ImageJ, and the 6 z-stacks of the images were subjected to maximum projection with maximum intensity. A rectangle was used to outline the entire perimeter of the image, and the fluorescence was measured. The area, and mean intensity of the image were measured.

### G. Statistics

Statistical analyses were performed using GraphPad Prism 8 (GraphPad software). Data are expressed as mean +/- SEM. Data from two groups were compared using two-tailed Unpaired Student’s t test. Multiple group comparisons were conducted using one-way ANOVA, two-way ANOVA, or RM Two-way ANOVA with post hoc tests as described in the appropriate figure legend. Statistical analysis was performed with an α level of 0.05. P values <0.05 were considered significant.

## Data availability

All source data supporting the findings are available within the article and Supplementary data. No custom scripts were generated for data analysis. Custom pipelines for data analysis are available from the corresponding author upon reasonable request.

## Acknowledgements

We thank Elyse Brozost and Rachell Rivera for technical assistance, Dr. Miho Nakajima and Dr. Nathaniel Heintz for the OTR Cre ON66 BAC transgenic mice, and Dr. Michael Gambello for the floxed Tsc2 mouse strain. We are grateful to all members of the Shrestha laboratory for critical feedback and discussions. Illustrations and schematic diagrams presented in Figures 1, 2, 3, 4 and Supplemental Figure 1 were generated using BioRender software (www.biorender.com). This study was supported by NIMH grant R01MH132795 and Sloan research fellowship (FG-2022-94485) to P.S, and NIDCD grant R01DC013073 to D.A.L. AVW is supported by Frances Velay Women and Science Research Fellowship and Simons STEM Scholar program at Stony Brook University. SL is supported by Scholars in Biomedical Sciences Program (T32GM148331).

## Author contributions

Conceptualization: PS

Methodology: AVG, AVW, SL, HS, JI, MD, BL

Investigation: PS, AVG, AVW, SL, HS, JI, MD, BL Funding acquisition: PS, DL

Supervision: PS, DL Writing – original draft: PS

Writing – review & editing: AVG, AVW Competing interests: None

