## Supplementary Figures for "Safety Signals Enable Single-Episode Active Avoidance paradigm and Expose Threat Generalization in Tuberous Sclerosis Complex"

### Supplementary Information

#### Supplementary Figure 1.

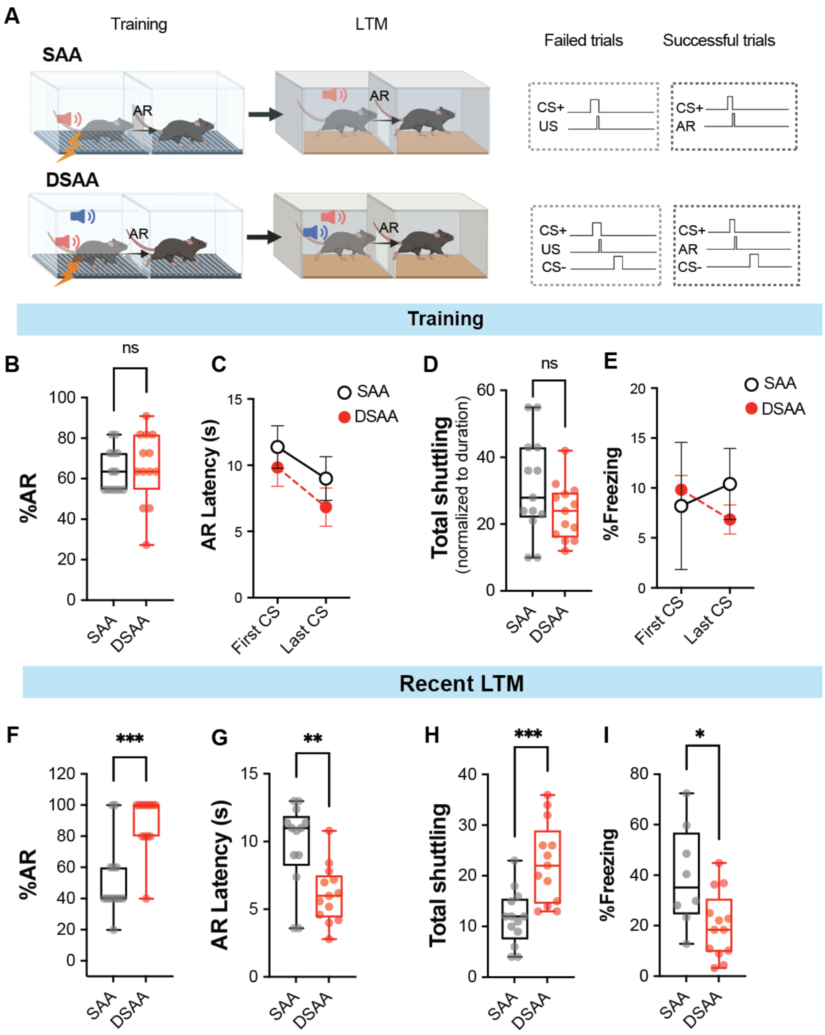


**A)** Schematic comparison of conventional signaled active avoidance (SAA) and the differential signaled active avoidance (DSAA) paradigm. In SAA, animals learn an instrumental response to a threat-predictive cue (CS+) to avoid the unconditioned stimulus (US), without presentation of a safety cue. In contrast, DSAA introduces a behaviorally non-contingent neutral cue (CS-), requiring animals to discriminate between threat (CS+) and safety (CS-). In both paradigms, failed trials result in US delivery immediately following CS+, whereas successful trials consist of an avoidance response (AR) during CS+, which terminates the cue and prevents the US. During DSAA training and long-term memory (LTM) testing, the CS− follows either the US (failed trials) or the AR (successful trials), respectively. **B)** Percentage of avoidance responses during training was comparable between SAA and DSAA groups. **C)** Reduction in avoidance latency from the first to the last CS+ presentation was similar across groups. **D)** Total shuttling behavior (normalized to session duration) and **E)** CS+-evoked freezing during training did not differ between paradigms. **F)** During recent LTM testing, DSAA animals exhibited significantly higher avoidance rates relative to SAA, indicating enhanced consolidation and retrieval. **G)** CS+-evoked avoidance latencies were further reduced in the DSAA group, demonstrating faster execution of the avoidance response. **H)** Total shuttling (normalized to duration) was increased in DSAA animals. **I)** Freezing responses remained prominent in SAA but were significantly attenuated in DSAA. **Statistical tests:** B, D, and F–I) Unpaired t-tests; C and E) RM two-way ANOVA. **Sample size:** n= 8-13 mice per group. *$p < 0.05$, **$p < 0.01$, ***$p < 0.001$; ns, not significant.

**Supplementary Figure 2.**

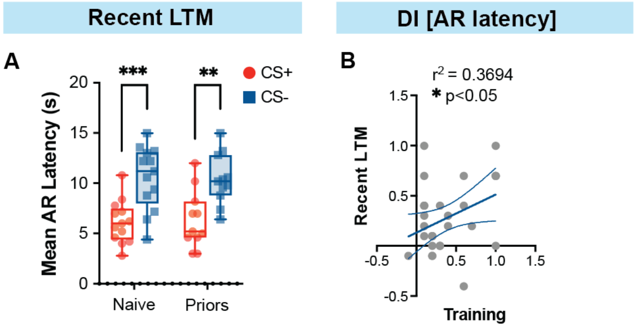


**A)** Both naïve and prior-conditioning groups exhibited appropriate cue-specific discrimination, reflected by significantly shorter avoidance latencies to the CS+ relative to the CS−. **B)** Discrimination indices for cue-specific avoidance latency during recent long-term memory (LTM) testing were positively correlated with discrimination indices measured during training, indicating that training-phase discrimination predicts subsequent memory precision (Pearson correlation: $R^2 = 0.3694$; $Y = 0.3793X + 0.1328$). **Statistical tests:** A) Two-way ANOVA with Bonferroni post hoc comparisons. B) Pearson correlation analysis. **Sample sizes:** A) n = 11–13 mice per group; B) n = 29 mice. *$p < 0.05$, **$p < 0.01$, ***$p < 0.001$.

**Supplementary Figure 3.**

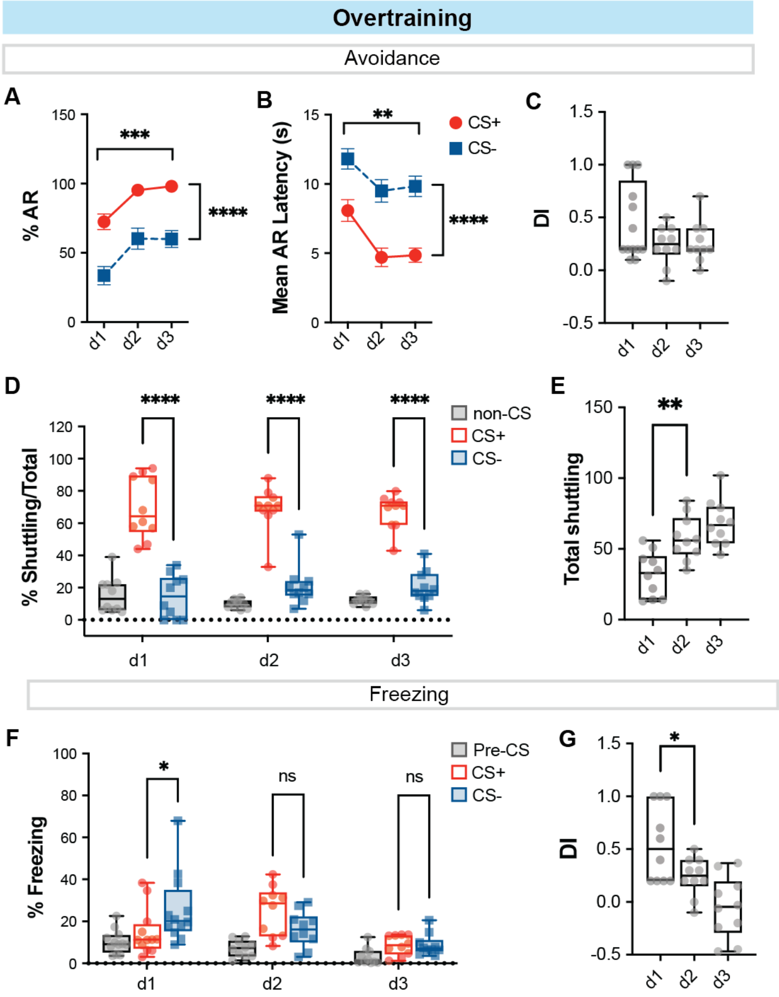

**A)** Following two additional days of overtraining, naïve DSAA mice exhibited enhanced avoidance performance to both CS+ and CS−. **B)** Mean avoidance response (AR) latencies progressively decreased across training days for both auditory cues. **C)** Box-and-whisker plots demonstrate that the discrimination index for avoidance latency remained stable across the three training sessions. **D)** The distribution of shuttling behavior (expressed as percentage of total shuttling) across CS+, CS−, and non-CS epochs remained consistent over training. **E)** Total shuttling increased from day 1 to day 2 and plateaued from day 2 to day 3. **F)** A differential freezing response to the CS− was evident on the first day of DSAA training; however, freezing declined to baseline levels on days 2 and 3. **G)** The freezing discrimination index significantly decreased from day 1 to day 2, consistent with reduced cue-specific freezing over time. **Statistical tests:** A, B) RM two-way ANOVA with Bonferroni post hoc test; C) one-way ANOVA; D, F) two-way ANOVA with Bonferroni post hoc comparisons; E, G) one-way ANOVA with Bonferroni post hoc comparisons. **Sample size:** n = 10 mice per group. *p < 0.05, **p < 0.01, ***p < 0.0001; ns, not significant.

**Supplementary Figure 4.**

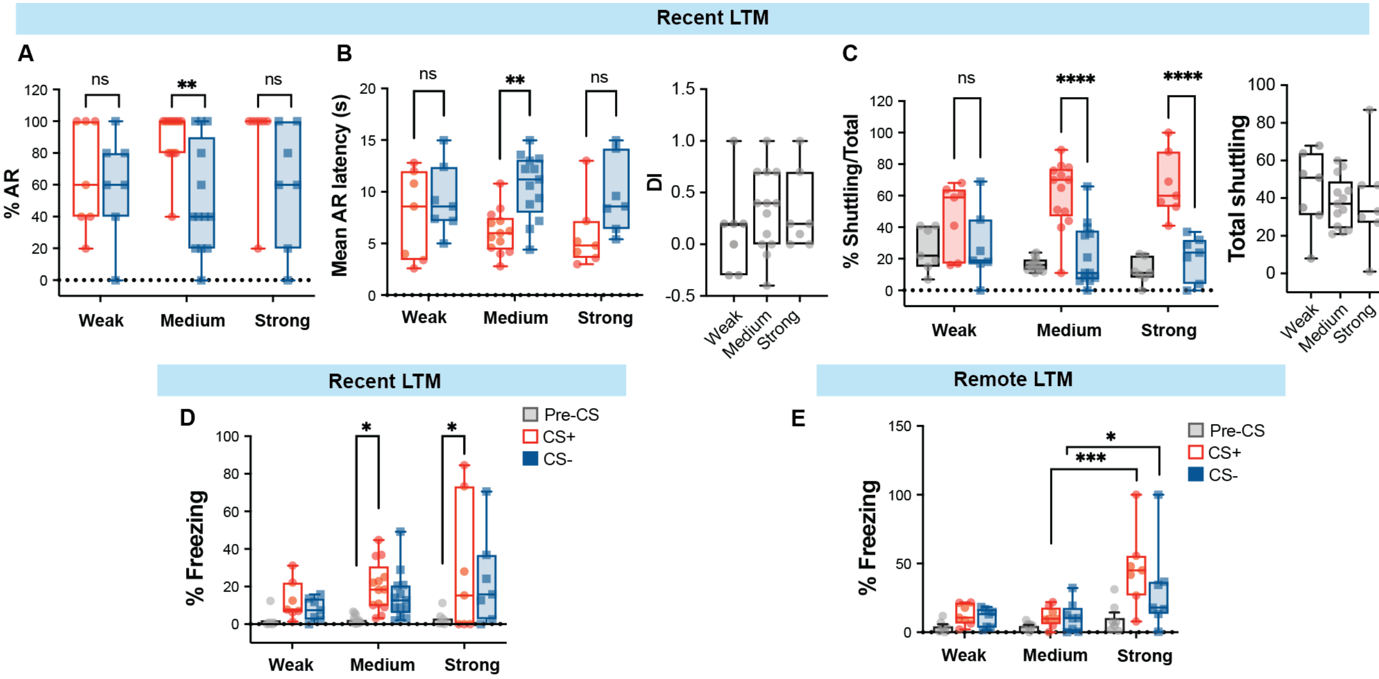


**A**) Box-and-whisker plots depicting differential avoidance responses (AR) in weak, medium, and strong US groups during the recent long-term memory (LTM) test. The medium US group exhibited significantly higher ARs to the CS+ compared to the CS−, indicating preserved discrimination. In contrast, both weak and strong US groups displayed generalized responding across cues. **B)** Left: Mean AR latency revealed cue discrimination in the medium US group, whereas latency responses were generalized in the weak and strong US groups. Right: The discrimination index did not differ significantly among the three groups. **C)** Left: Shuttling distribution demonstrated appropriate divergence toward the CS+ relative to the CS− and non-CS epochs in the medium and strong US groups; this divergence was absent in the weak US group. Right: Total shuttling did not differ across groups. **D)** During the recent LTM test, freezing responses were higher to the CS+ than the CS−, with group-dependent differences in CS−-evoked freezing. **E)** At remote LTM, the strong US group showed significantly elevated freezing to both CS+ and CS− compared to the weak and medium US groups, indicating enhanced generalized defensive responding. **Statistical tests:** A, B-left, C-left, D, E) Two-way ANOVA with Bonferroni post hoc tests; B-right, C-right) One-way ANOVA. Sample size: n = 7–13 mice per group. p < 0.05, p < 0.01, p < 0.001, p < 0.0001; ns, not significant.

**Supplementary Figure 5.**

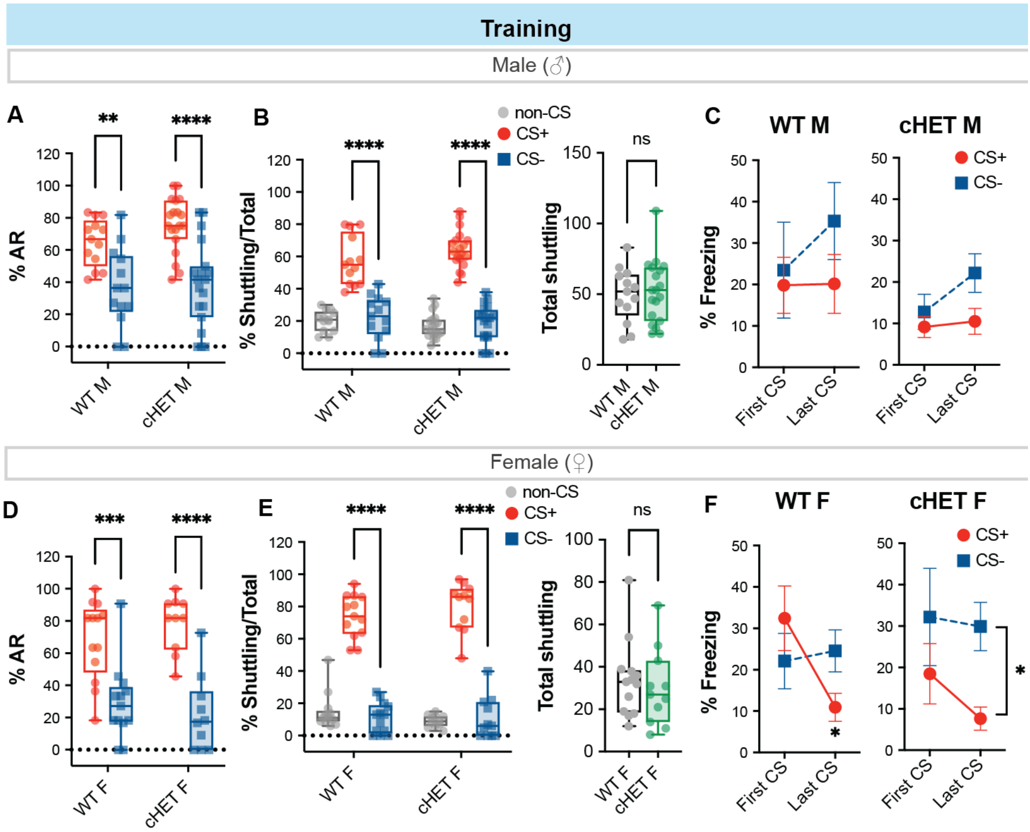


**A)** Box-and-whisker plots showing differential avoidance responses (AR) in wildtype (WT) and OTR.Tsc2$^{f/+}$ (cHET) male mice during DSAA training. **B)** Left: Shuttling distribution revealed appropriate divergence toward the CS+ relative to the CS− and non-CS epochs in both WT and cHET males. Right: Total shuttling did not differ between genotypes. **C)** Percent freezing during DSAA training in WT and cHET male mice, quantified during the Pre-CS baseline, first CS, and last CS epochs. **D)** Box-and-whisker plots depicting differential avoidance responses (AR) in WT and OTR.Tsc2$^{f/+}$ (cHET) female mice during DSAA training. **E)** Left: Shuttling distribution showed appropriate CS+-biased divergence relative to CS− and non-CS epochs in both WT and cHET females. Right: Total shuttling did not differ between genotypes. **F)** Percent freezing during DSAA training in WT and cHET female mice, measured during the Pre-CS baseline, first CS, and last CS epochs. **Statistical tests**: A, B-left, D, E-left) Two-way ANOVA with Bonferroni post hoc tests; B-right, E-right) Unpaired t-test; C, F) RM Two-way ANOVA with Bonferroni post hoc test. Sample size: n = 9–19 mice per group. *p<0.05, **p<0.01, ***p<0.001, ****p<0.0001, ns not significant.

**Supplementary Figure 6.**

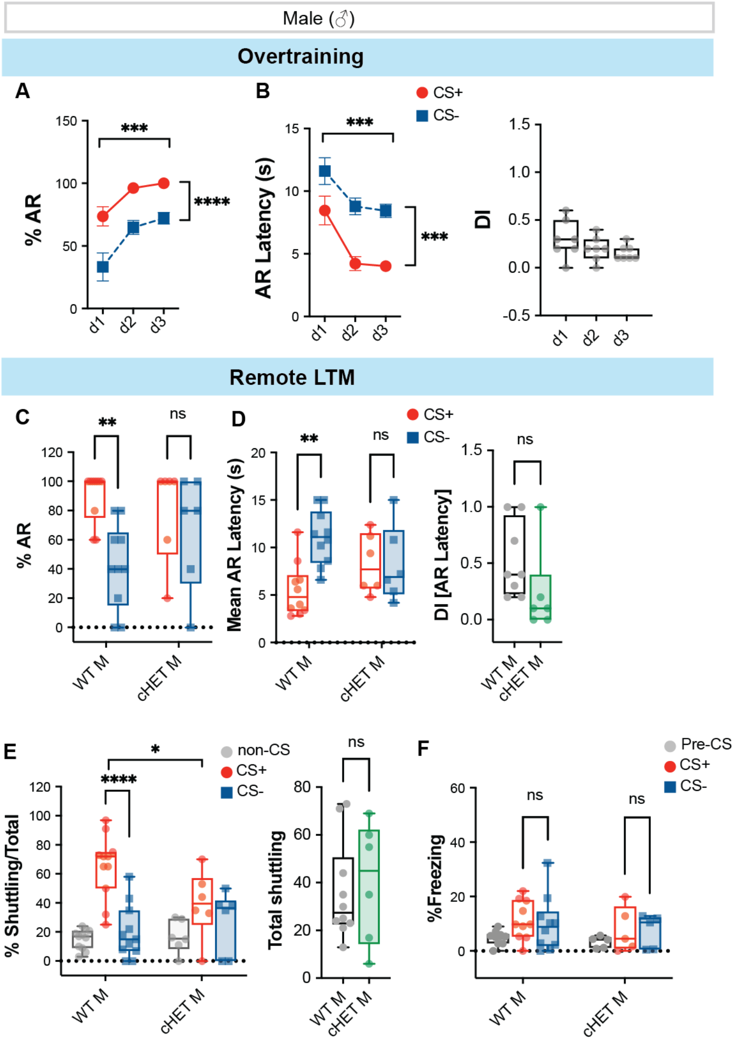


**A)** XY plots illustrating the learning curve for avoidance responses to CS+ and CS− across successive days of training (overtraining) in OTR.Tsc2^f/+^ (cHET) male mice. **B)** Left: Avoidance latency progressively decreased over additional training days in cHET males, indicating improved task performance. Right: The discrimination index remained stable across all three training days. **C)** During remote long-term memory (LTM) testing, overtrained wildtype (WT) males exhibited clear differential avoidance between CS+ and CS−, whereas overtrained cHET males displayed generalized responding across cues. **D)** Mean avoidance latency at remote LTM reflected preserved cue discrimination in WT males but generalized latency responses in cHET males. **E)** Shuttling distribution demonstrated appropriate CS+-biased divergence relative to CS− and non-CS epochs in overtrained WT males; in contrast, distributions converged in cHET males despite comparable total shuttling between genotypes. **F)** Cue-evoked freezing remained minimal for both CS+ and CS− during remote LTM in overtrained WT and cHET males. **Statistical tests:** A, B-Left) RM Two-way ANOVA with Bonferroni post hoc test; B-Right) One way ANOVA; C, D-Left, E-Left, F) Two-way ANOVA with Bonferroni post hoc test; D-Right, E-Right) Unpaired t-test. Sample size: n = 5-10 mice per group. *p<0.05, **p<0.01, ***p<0.001, ****p<0.0001, ns not significant.
